# Complex eco-evolutionary dynamics induced by the coevolution of predator-prey movement strategies

**DOI:** 10.1101/2020.12.14.422657

**Authors:** Christoph Netz, Hanno Hildenbrandt, Franz J. Weissing

## Abstract

The coevolution of predators and prey has been the subject of much empirical and theoretical research, which has produced intriguing insights into the intricacies of eco-evolutionary dynamics. Mechanistically detailed models are rare, however, because the simultaneous consideration of individual-level behaviour (on which natural selection is acting) and the resulting ecological patterns is challenging and typically prevents mathematical analysis. Here we present an individual-based simulation model for the coevolution of predators and prey on a fine-grained resource landscape. Throughout their lifetime, predators and prey make repeated movement decisions on the basis of heritable and evolvable movement strategies. We show that these strategies evolve rapidly, inducing diverse ecological patterns like spiral waves and static spots. Transitions between these patterns occur frequently, induced by coevolution rather than by external events. Regularly, evolution leads to the emergence and stable coexistence of qualitatively different movement strategies. Although the strategy space considered is continuous, we often observe discrete variation. Accordingly, our model includes features of both population genetic and quantitative genetic approaches to coevolution. The model demonstrates that the inclusion of a richer ecological structure and higher number of evolutionary degrees of freedom results in even richer eco-evolutionary dynamics than anticipated previously.

## 1. Introduction

Predator-prey coevolution has fascinated biologists for decades (Cott 1940, Pimentel 1961, Levin & Udovic 1977, Dawkins & Krebs 1979). In the ecological arena, predator-prey interactions can lead to complex non-equilibrium dynamics (Turchin 2003). On top of the ecological interactions, an evolutionary arms race is taking place where adaptive changes in one species impose new selective pressures on the other species. Experimental findings suggest that the ecological and the evolutionary dynamics can be intertwined in an intricate manner (Yoshida et al. 2003, 2007, Becks et al. 2010). In natural systems, it is a major challenge to unravel this complexity (Hendry 2019). In view of all this, it is no surprise that theoretical models have played a crucial role for understanding the ecology and evolution of predator-prey interactions (Fussmann et al. 2007, Govaert et al. 2019).

Predator-prey coevolution models have traditionally been based on the frameworks of population genetics (Nuismer et al. 2005, Kopp & Gavrilets 2006, Cortez & Weitz 2014, Yamamichi & Ellner 2016), quantitative genetics (Gavrilets 1997, Mougi & Iwasa 2010, Cortez 2018), and adaptive dynamics (Dieckmann & Law 1996, Marrow et al. 1996, Flaxman & Lou 2009). Each approach encompasses a diversity of models. The approaches differ in their assumptions on the nature of genetic variation (discrete vs continuous), the occurrence and distribution of mutations, and the interaction of alleles within and across loci, but they have in common that they strive for analytical tractability. To achieve this, virtually all analytical models make highly simplifying assumptions on the traits that are the target of selection. For example, predators are characterized by a one-dimensional attack strategy, prey by a one-dimensional avoidance strategy, and prey capture rates are assumed to be maximal when the predator attack strategy matches the prey avoidance strategy (Van der Laan & Hogeweg 1995, Dieckmann & Law 1996, Marrow et al. 1996, Gavrilets 1997, Nuismer et al. 2005, Kopp & Gavrilets 2006, Yamamichi & Ellner 2016, see Abrams 2000 for a general overview). Such simplification allows for an elegant and seemingly general characterisation of coevolutionary outcomes, but the question arises whether it captures the essence of predator-prey interactions, which in natural systems are mediated by complex behavioural action and reaction patterns.

Experiments have demonstrated that predators and prey show strong behavioural responses to each other (Gilliam & Fraser 1987, Savino & Stein 1989, Ehlinger 1990, Hammond et al. 2007, Simon et al. 2019). These responses differ across species and spatial scales, and they are likely the product of behavioural strategies that incorporate various information sources from the environment. In the literature, behavioural interactions between predators and prey are often discussed in game-theoretical terms as ‘behavioural response races’ (Sih 1984, 2005) or ‘predator-prey shell games’ (Mitchell & Lima 2002). These games usually play out in space, where resources, prey and predators are heterogeneously distributed among different patches. Prey have to balance foraging for a resource with predator avoidance, while predators are faced with the task of predicting prey behaviour. A full dynamical analysis of such interactions is a forbidding task (Flaxman & Lou 2009). Therefore, analytical approaches typically use short-cuts, such as the assumption that at all times individual predators or prey behave in such a way that they maximize their fitness under their given local circumstances (Gilliam & Fraser 1987, Abrams 2007). It is often doubtful whether such short-cuts are realistic. For example, it is unlikely that evolution will fine-tune behaviour to such an extent that it is locally optimal under all circumstances and when conditions rapidly change (McNamara & Houston 2009, McNamara & Weissing 2010). To make progress, we therefore have to consider the option of abandoning analytical tractability and instead adopt a simulation-based approach that can make more realistic assumptions on the ecological setting and the traits governing the behaviour of predators and prey.

Several such simulation models have been developed in the past (Huse et al. 1999, Kimbrell & Holt 2004, Flaxman et al. 2011, Patin et al. 2020). All these models are individual-based, meaning that both the ecology and the evolution of predator-prey interactions reflect the fate of individual agents. Such an approach is natural because selection acts on individual characteristics, while population-level phenomena are aggregates and/or emerging features of these characteristics. Additionally, individual-based models enable the implementation of considerable mechanistic complexity, both at the level of individual behaviour and with regard to the environmental setting. The existing individual-based models have shown that they can reproduce findings of analytical models (Huse et al. 1999, cf. Iwasa 1982) and theoretical expectations such as the ideal free distribution (Flaxman et al. 2011). As illustrated by the recent study of Patin and colleagues (2020), such models can unravel the importance of random movement, memory use, and other factors that are often neglected.

Still, current individual-based models for the coevolution of predators and prey are limited by the fact that individual interactions are considered at a coarse-grained spatial scale (for an exception see Kimbrell & Holt 2004). The environment is assumed to be structured in discrete patches and individuals have the task of choosing a patch that provides an optimal balance between resource abundance and safety. Within patches, predator-prey interactions are governed by patch-level population dynamics and not based on single individuals moving and behaving in space. Accordingly, these models do not capture interactions that take place at the individual level.

Here, we therefore consider a model where individuals move and interact in more fine-grained space, where only few individuals co-occur at the same location. Predators and prey evolve situation-dependent movement strategies that determine the likelihood of finding food resources and avoiding predation. On an ecological (or behavioural) timescale, individual predators and prey repeatedly take movement decisions based on an inherited strategy that evaluates the ‘suitability’ of environmental situations. We assume that predator and prey individuals continually scan their environment in various directions and sense the local density of resources, prey and predators in these directions. Based on these inputs, each individual judges the suitability of the various movement directions (including its current location), and makes a step in the direction of highest suitability. The movements individuals make determine their survival and foraging success in case of prey, and their prey capture rate in case of predators. These success rates in turn affect the number of offspring produced. On an evolutionary timescale, successful individuals transmit their movement strategy to many offspring, subject to some mutation. Over the generations, successful strategies will spread, thus potentially changing the nature of ecological interactions and the spatial pattern of resource densities and abundances.

On purpose, we have kept our model as simple as possible. We are more interested in obtaining conceptual insights than in accurately representing any specific real-world predator-prey system. To this end, we will ask questions such as: What kinds of predator-prey movement strategies do evolve and can they be classified in relation to each other? Which information sources are used by predators and prey, respectively? How do predator-prey interactions reflect and shape the resource landscape? Are the predictions of earlier models such as regular trait cycles recovered in our model? Are the evolved populations monomorphic, or do different prey and predator strategies coexist in the same population? How predictable is the outcome of coevolution; do replicate simulation runs produce qualitatively similar outcomes? How fast is evolutionary change in relation to ecological change; does the interplay of ecology and evolution lead to novel eco-evolutionary patterns and dynamics?

## 2. The Model

We consider an individual-based model for the antagonistic coevolution of movement strategies between a population of herbivores (prey) and a population of predators in a spatially explicit environment.

### Ecological Interactions

To be concrete, we imagine the prey to be herbivores feeding on a resource, henceforth called grass, and predators feeding on herbivores. All individuals live in an environment consisting of a grid of 512*512 cells with wrapped boundaries. A cell can host one or several herbivores and predators, but this is unlikely since the number of grid cells is an order of magnitude larger than the number of individuals. Ecological interactions occur in discrete time steps. A time step (we think of a day) contains a grass growth phase, a movement phase and a foraging phase with predator-prey interactions. Grass grows at a constant rate of 0.01 per time step up to a maximum level of 1. Next, herbivores and predators move between cells based on their inherited movement strategy as described below. Herbivores visiting a cell deplete the grass and gain the corresponding amount of energy. Herbivores occupying the same cell share the amount of grass on that cell. If a predator encounters a herbivore on the same cell, the predator succeeds to capture the herbivore with probability 0.5, in which case the herbivore is killed and the predator gains one unit of energy. If several predators co-occur with several herbivores (which only rarely ever happens), the successful predators kill and consume all the herbivores present in the cell. The killed herbivores are equally distributed between successful predators.

In the graphs below, the ‘performance’ of the predator and herbivore population is quantified as follows: Predator performance is equal to the average lifetime consumption of prey items per predator. Herbivore performance corresponds to the average lifetime resource intake per herbivore, where lifetime intake was set to zero for those herbivores that did not survive predation.

### Movement strategies

Movement strategies are based on the evaluation of nearby cells (see fig. 1). For each evaluated grid cell, a ‘suitability score’ *S* is calculated. *S* is the weighted sum *S* = *w_g_G* + *w_h_H* + *w_p_P*, where *G*, *H* and *P* are the grass density, herbivore density, and predator density in the cell, respectively. The weighing factors *w_g_*, *w_h_*, and *w_p_* are genetically encoded and transmitted from parent to offspring.

**Figure 1:**
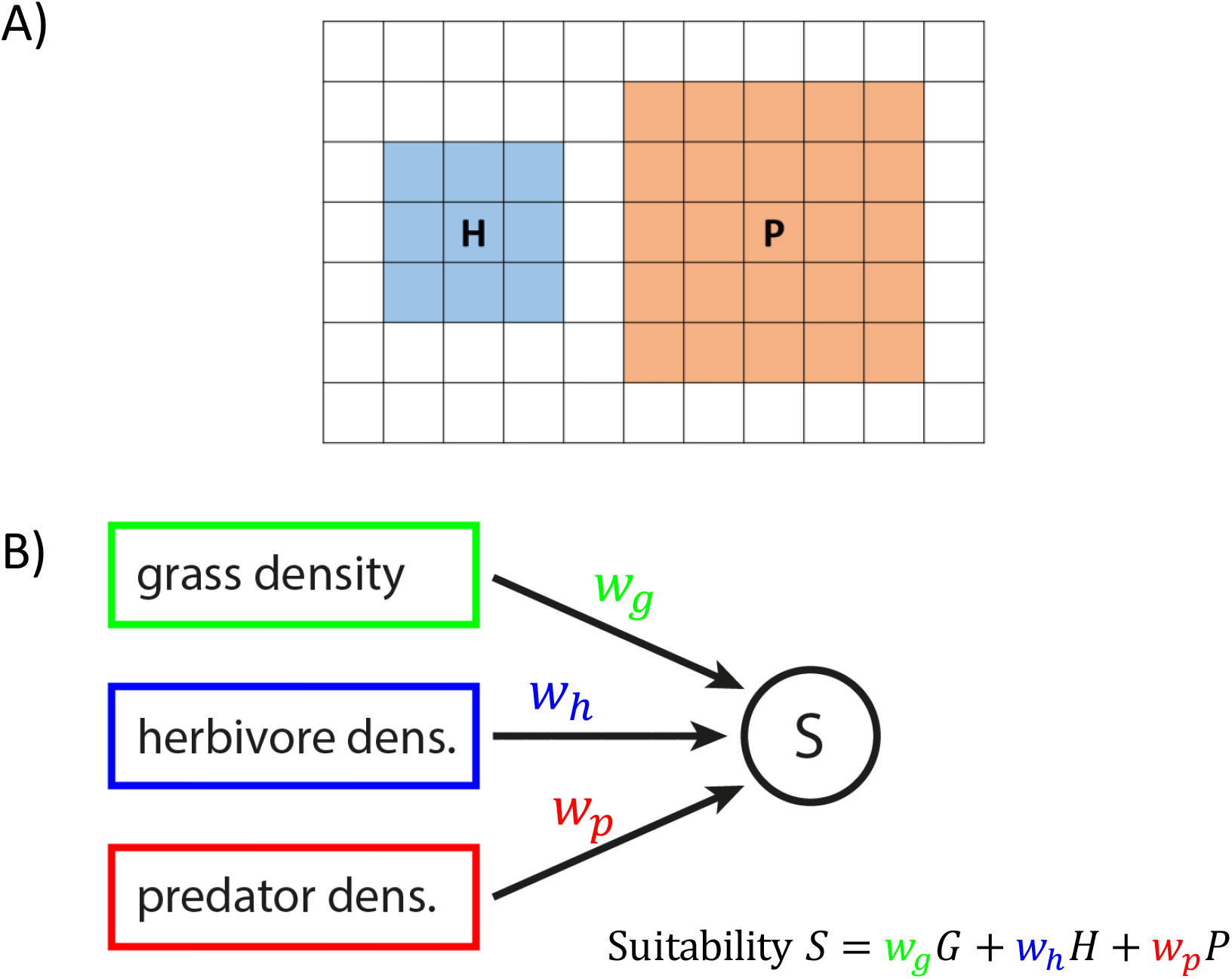
Movement and decision-making. A) Herbivores and predators move on the same rectangular grid, but their movement range per time step can be different. The plot illustrates the movement range of herbivores (blue, radius = 1) and predators (red, radius = 2) for our standard configuration. B) Individuals evaluate all cells in their movement range as to their ‘suitability’ and move to the cell of highest suitability. The suitability *S* of each cell is the weighted sum of the local grass density *G*, the local herbivore density *H*, and the predator density *P*, where the weighing factors *w_g_*, *w_h_*, and *w_p_* are genetically determined and hence evolvable.

Grass density represents the total amount of grass in a given cell, while for herbivore and predator densities we convoluted the otherwise discrete presence-absence values of agents via a Gaussian filter (Lindeberg 1994). This yields continuous values of herbivore and predator densities that are diffused around the actual positions of the agents, much like we would expect from olfactory or similar cues. Individuals evaluate all cells in their movement range, and move to the cell with the highest suitability. In the simulations shown, the movement range of herbivores had a radius of one, and the predators a radius of two (9 cells and 25 cells, fig. 1a). If predators have the same movement radius as their prey, herbivores can reliably escape predation by moving away from high predator densities. Only when predators can move further than herbivores, both parties need to predict the behaviour of the other party, and interactions become more intricate.

### Evolution of the evaluation mechanism

We consider haploid parthenogenetic populations with discrete, non-overlapping generations. The population sizes are constant at the start of each generation. The herbivore population is initiated each generation to 10,000 individuals and is successively diminished by predation. The predator population size remains constant (either 10,000 or 1,000 individuals). At the start of a new generation, individuals are placed at random in the ‘dispersal range’ around the position of their parent. In the simulations shown below we used a dispersal radius of 1 or 10 (for both species).

Each individual has three gene loci with alleles *w_g_*, *w_h_*, and *w_p_* that correspond to the weights encoding the evaluation of environmental suitability and, hence, determine the individual’s movement strategy. The movement strategy determines the types of habitat most likely visited and therefore the individual’s intake rate and, in the case of the herbivore, the individual’s probability of escaping predation. At the end of a generation (after 100 time steps), surviving individuals produce offspring where the number of offspring is proportional to their total intake. Each offspring inherits the genetic parameters of its parent, subject to rare mutations. A mutation occurs with probability μ=0.001 per locus, in which case the original value is changed by an amount drawn from a Cauchy distribution with location 0 and scale parameter 0.001.

## 3. Results

### Transitions between spatial patterns

Figure 2 shows a representative simulation run of our model for the situation that offspring dispersal is local (dispersal radius 1) and predators and herbivores occur in equal proportions. Within generations, the movement strategies of predators and herbivores induce ecological patterns that are characteristic for these strategies (fig. 2B). Over generations, the movement strategies evolve; the evolutionary dynamics of the weights are shown in Supplementary figure S1. The change in movement strategies alters the ecological pattern. Even when averaged over the landscape, the performance of predators and herbivores neither reaches an equilibrium nor stable oscillations (fig. 2A). Instead, evolution is stochastic, highly dynamic and continually produces new strategies and ecological patterns. The performance of predators and herbivores undergoes distinct phases. In the first half of the simulation (fig. 2A), we see about 8 phases with a relatively stable configuration, between which the system rapidly shifts. Each of these configurations corresponds to a specific combination of movement strategies (Supplementary fig. S1) and an associated ecological pattern (fig. 2B.1-4). Predators and herbivores can, for example, be distributed relatively homogeneously (fig. 2B.2) or in static spatial clusters forming spot patterns (fig. 2B.3); they can also produce dynamic spatial patterns such as rotating spirals (fig. 2B.4).

**Figure 2:**
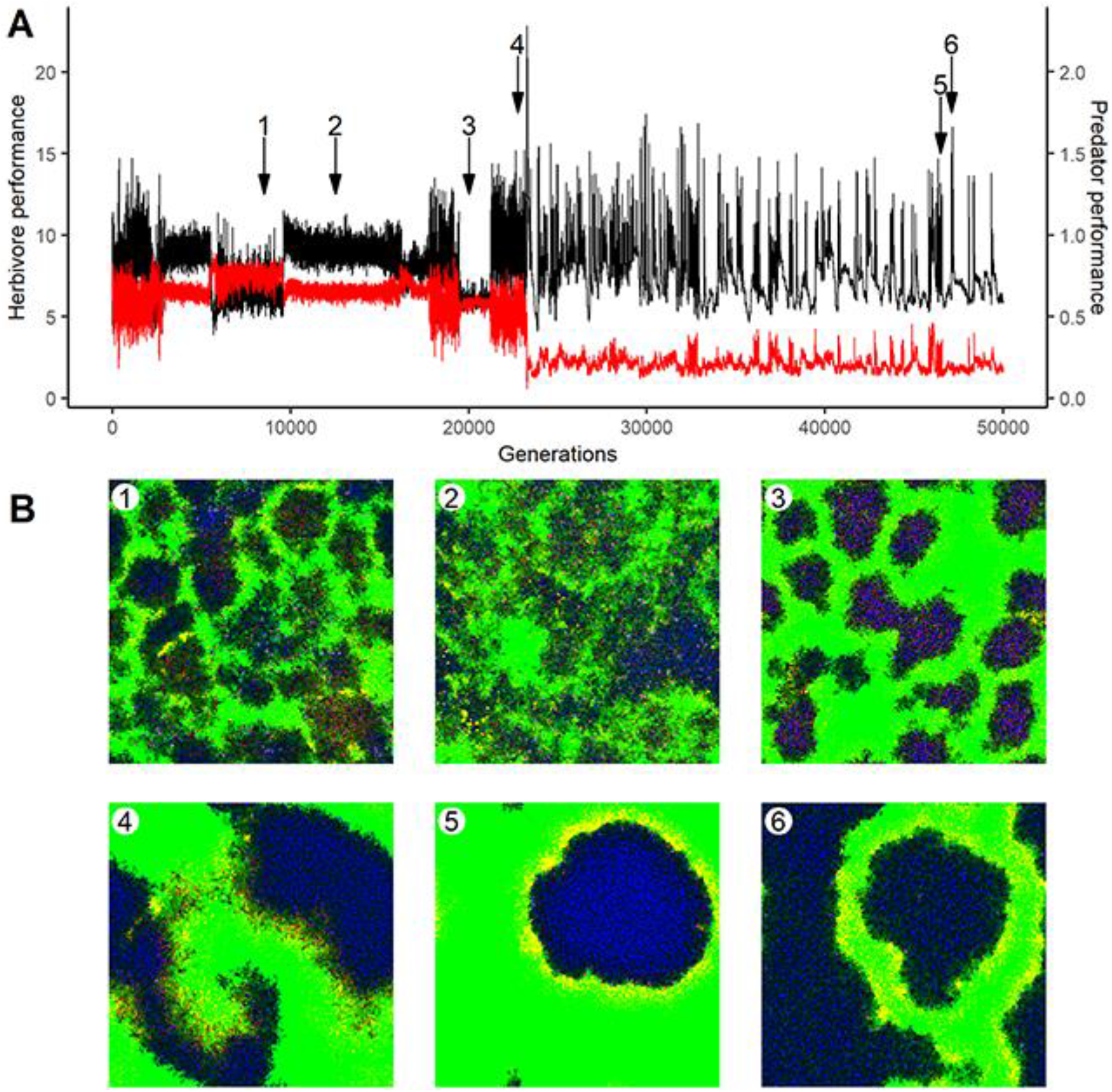
Eco-evolutionary pattern formation. (A) Change in the performance of predators (red, average number of consumed prey per predator) and herbivores (black, average lifetime consumption of resources per herbivore, set to zero for individuals captured by predators) over 50,000 generations. (B) Landscape snapshots at six time points of the evolutionary trajectory (indicated by arrows in panel A). Grass density is shown in green, herbivore density in blue and predator density in red. Other colors emerge from additive color mixing, yellow for example signifies areas of high grass and predator density, purple that herbivores and predators occupy the same area. Predator-prey ratio 1:1; offspring dispersal radius 1.

In the second half of the simulation, the stable configurations disappear and the system starts to exhibit rapid large-scale fluctuations. This phase is characterized by large herbivore aggregations enclosed by predators that remain stable for multiple generations (fig. 2B.5). Repeatedly, the emergence of new movement strategies allows the herbivores to break out of these aggregations and spread rapidly across the landscape (fig. 2B.6), until a new stable spatial formation between predators and herbivores is reached. The pattern in figure 2B.5 corresponds to a period of low herbivore performance, while herbivore performance is high in a pattern like figure 2B.6.

The performance phases and the corresponding ecological patterns reflect the evolved movement strategies of herbivores and predators. For example, dynamic spatial patterns such as spirals tend to be produced by herbivores that move into areas of high grass density, while predators chase after them. In contrast, spatial aggregations as in figure 2B.5 reflect a ‘sit-and-wait’ strategy of the predator. Individual predators prefer high grass densities and tend to remain there, while herbivores avoid predators; the predators can thus monopolize high grass patches and consume those herbivores that are eventually lured in by the high grass density. Although simulation replicates can vary a lot in their behaviour (Supplementary material videos 1-3), the simulation run featured above is characteristic for simulations with local offspring dispersal and a 1:1 predator-prey ratio. Under global offspring dispersal, a landscape structure cannot emerge in the timeframe of our simulations (100 movement steps per generation); under these conditions, the herbivore population often went practically extinct.

### Polymorphism and trait cycles

We now turn to the case of a 1:10 predator-prey ratio and intermediate dispersal radius (r=10). Figure 3 shows the change of herbivore and predator performance over time and the evolution of weighing factor *w_p_* in the herbivore population (for the evolution of all six weights see Supplementary fig. S2). Under these conditions, spatial patterns are less pronounced and spatial differentiation is more limited (see Supplementary fig. S3, Supplementary Material video 4). As shown in the top panel of figure 3, the overall performance level of both populations is subject to relatively regular oscillations. In all but one of the simulation replicates we investigated (n=30), a pronounced polymorphism in the herbivore strategy evolved (fig. 3B): the evolved weight *w_p_* is strongly negative in some herbivores (purple) and positive in all others (green). Since *w_p_* determines how the density of predators affects the suitability assessment of a location by a herbivore individual, the two morphs respond very differently to the presence of predators. The purple morph (with *w_p_* < 0) avoids locations with a high density of predators, as one would expect. In contrast, the green morph is attracted by locations with a high predator density. However, the avoidance of the purple morph is much stronger than the green morph’s preference, and the two morphs differ more generally in their movement strategy: The green morph moves primarily according to grass densities, while the purple morph increasingly moves to avoid predator densities (see information usage, Supplementary fig. S4).

**Figure 3:**
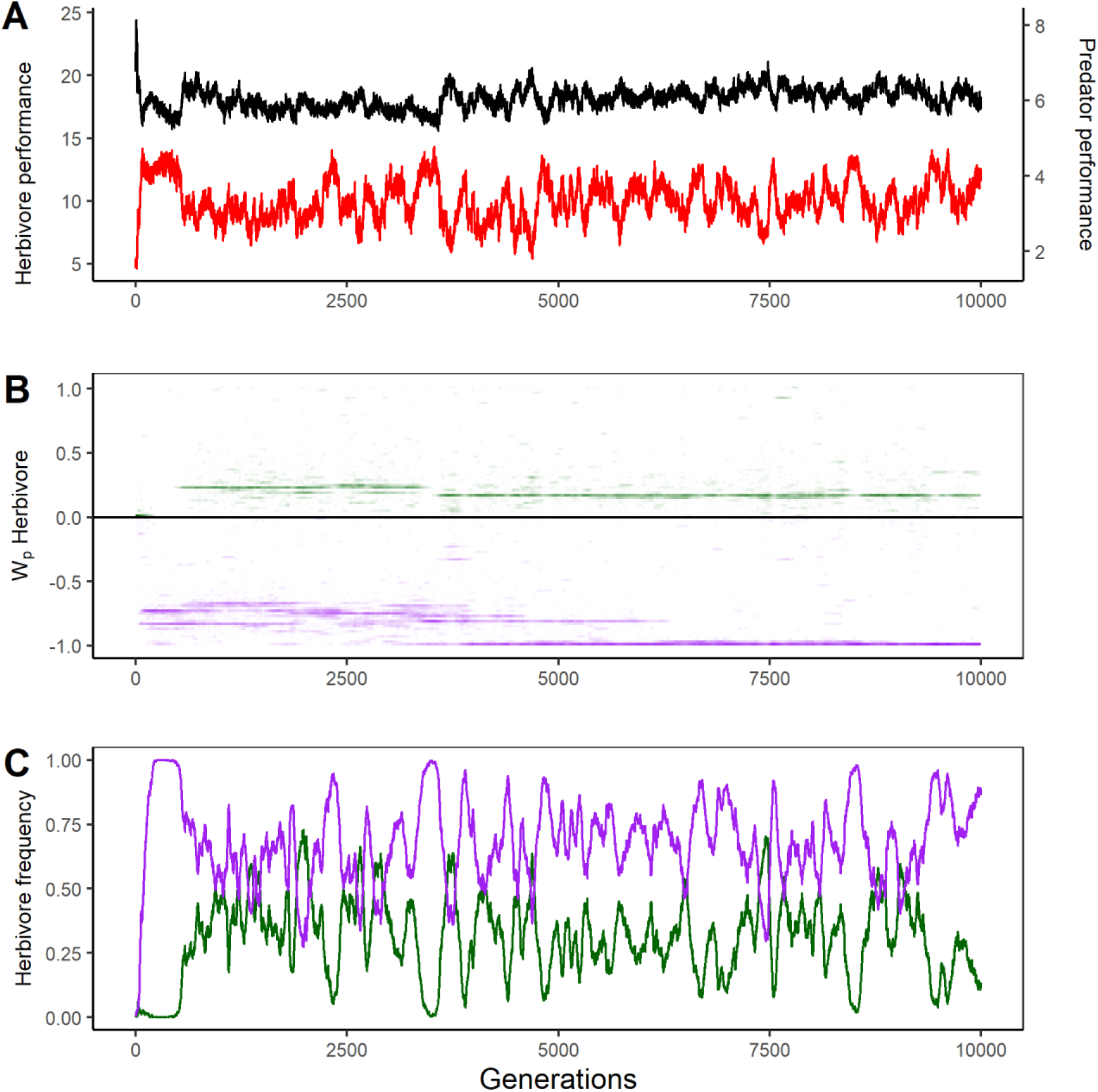
Polymorphism and trait cycles. A) Change in the performance of predators (red, average number of consumed prey per predator) and herbivores (black, average lifetime consumption of resources per herbivore, set to zero for individuals captured by predators) over 10,000 generations. B) Evolution of the distribution of the weighing factor *w_p_* (= the weight given to predator density) in the herbivore population. In all simulations, a polymorphism emerged, consisting of a predator-avoiding morph (purple) and a predator-prone morph (green). To enhance the visibility of rare morphs, weight values were multiplied with 100 and tanh-transformed, while weight frequencies were square-root transformed. C) Persistent oscillations in the relative frequency of the predator-avoiding (purple) and the predator-prone (green) morph. Predator-prey ratio 1:10; dispersal radius r=10.

At first sight, the behaviour of the green morph seems counterintuitive, but can be explained by the fact that the predators evolved an aversion against locations with a high predator density (see Supplementary fig. S2), presumably in order to avoid competition with other predators. This means that predators tend to move away from locations with high predator density, leaving these locations predator-free in the next time step. As long as predators are self-avoiding, the green morph is therefore a viable strategy. Indeed, the green morph suffers less from predation than the purple morph. In contrast, the purple morph is more efficient in exploiting resources. Hence the polymorphism reflects two contrasting strategies when facing a trade-off between predator avoidance and resource attraction.

As shown in figure 3C, the two morphs exhibit stochastic but regular oscillations in relative frequency. Overall, the predator-avoiding purple morph is more frequent, but there are periods where the predator-prone green morph has the highest frequency. The comparison of figures 3A and 3C reveals that the fluctuations in herbivore and predator performance are driven by the oscillation in morph frequency. In particular, predator performance (= overall predation success) is strongly correlated with the frequency of the purple morph, which is less able to avoid predation than the green morph.

The described polymorphism is a characteristic feature of all simulations with local offspring dispersal (dispersal radius 1 and 10); it did not occur in simulations with global offspring dispersal.

### Evolutionary oscillations

When considering other simulation settings, we observed a variety of additional phenomena. For example, we considered the implications of limiting the available information to one or both parties. To illustrate this, we here show a simulation, in which all three information channels are available to herbivores, while predators can only sense the density of other predators (Figure 4). In this scenario, the weighing factor *w_p_* does not only exhibit a pronounced polymorphism in in the herbivores but also in the predator population (Fig. 4B and C; the evolution of the other weights is shown in Supplementary Fig. S5). At the start of the trajectory shown (at generation 9,000), both predator and herbivore populations have a morph with a positive preference for predator density and a morph with a negative preference for predator density. Each of these morphs exploits a strategic weakness in one of the other morphs. The predator-avoiding (purple) herbivore morph is adapted to the predator-prone (red) predator morph and will almost never be caught by such predators. The red predator morph exploits the predator-attracted (green) herbivore morph, this morph in turn profits from the predator-avoiding (blue) predator morph by walking in its steps. The blue predator morph in turn is the most efficient predator of the equally predator-avoiding purple herbivore morph. As long as all four morphs are present (between generations 9,000 and 10,500), they oscillate rapidly and in a regular fashion.

**Figure 4:**
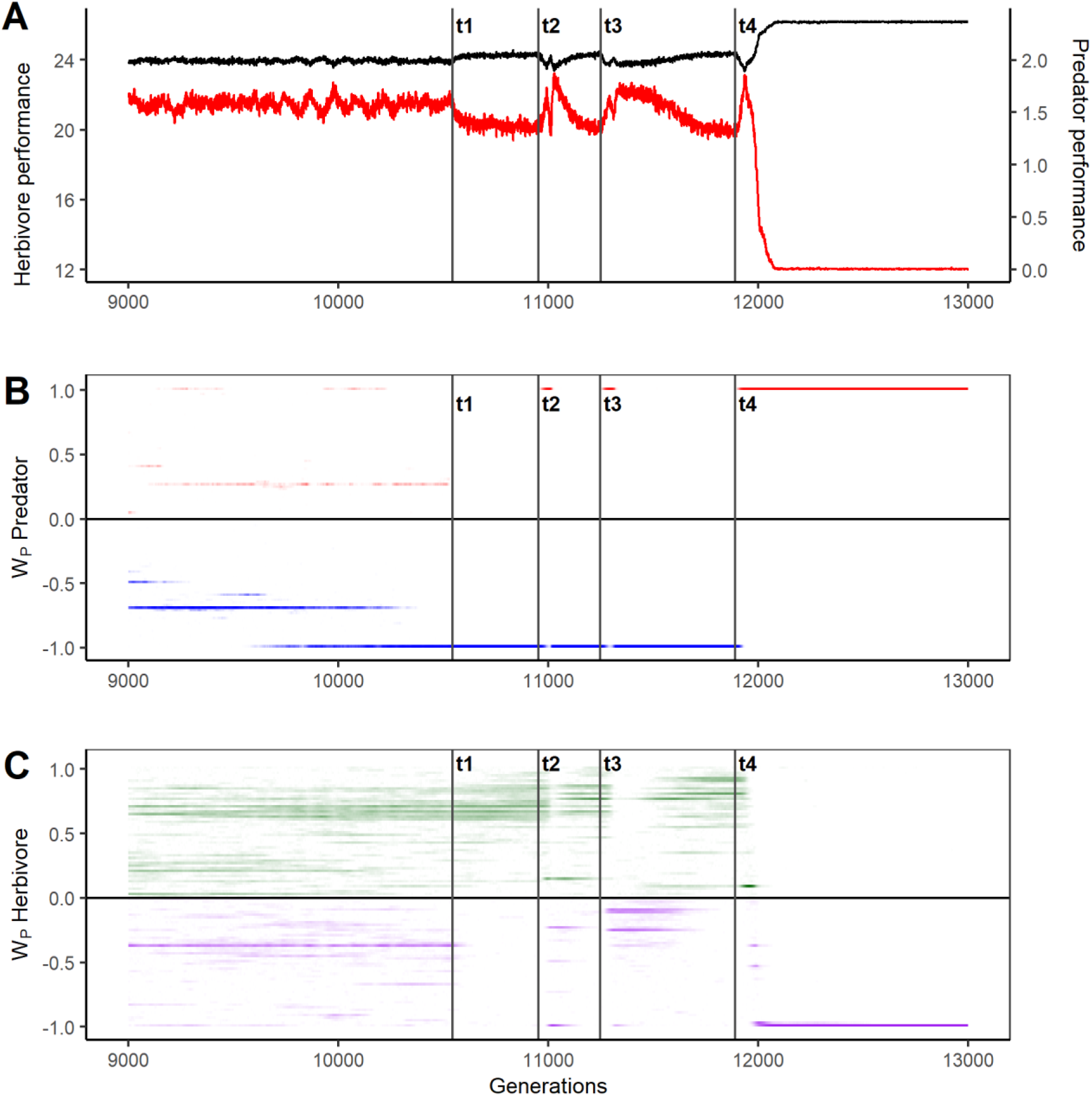
Predator extinction and two types of trait cycles. Cut-out from a simulation in which predators could only sense the density of other predators. A) Change in the performance of predators (red, average number of consumed prey per predator) and herbivores (black, average lifetime consumption of resources per herbivore, set to zero for individuals captured by predators). B) Evolution of the distribution of the weighing factor *w_p_* (= the weight given to predator density) in the predator population. Two distinct predator morphs emerge, a predator-prone morph (red) and a predator-avoiding morph (blue). C) Evolution of the distribution of the weighing factor *w_p_* (= the weight given to predator density) in the herbivore population. Two distinct herbivore morphs emerge, a predator-prone morph (green) and a predator-avoiding morph (purple). To enhance the visibility of rare morphs, weight values in B) and C) were multiplied with 100 and tanh-transformed, while weight frequencies were square-root transformed. Predator-prey ratio 1:10; dispersal radius r=10.

Around generation 10,500, the system enters a new phase, when the red predator morph goes extinct (at t1, see fig. 4). The purple herbivore morph, being specialized on the red predator morph, follows suit and also goes extinct. For about 400 generations, the performance of the predator population (= the overall catching rate of prey) fluctuates at a relatively low level. This changes with the arrival of a beneficial mutation in the predator population (at t2, fig. 4): a variant of the red (predator-prone) morph suddenly reappears and becomes very frequent, exploiting the abundant green herbivore morph. Here, the predator performance has a first peak. At this point the purple herbivore morph is reintroduced via mutation and becomes very frequent. As the purple morph increases, the red predator morph becomes rare and finally goes extinct again. The blue predator morph, which remained present in the population, takes over again and benefits from the common purple herbivore morph. Here, the predator performance peaks a second time. As the predator population is again monomorphic and blue, the green herbivore morph becomes more common and the purple herbivore morph goes extinct; the system returns to its original state. This cycle happens twice-over (t2, t3, fig. 4), but the third time the red morph reappears (at t4, fig. 4), the blue predator morph goes extinct. As the purple herbivore morph rapidly takes over the population, the now monomorphic predators do not catch prey anymore and their predation rate goes to practically zero. The population also cannot shift back to a negative preference, because this morph has gone extinct, and a new adaptive mutation does not arise in time. Hence the predators practically go extinct.

## 4. Discussion

We here presented a model considering the antagonistic coevolution of movement strategies. Predators and prey move around in a fine-grained resource landscape where they encounter antagonists and conspecifics. Movement strategies are based on three heritable parameters determining how individuals assess the densities of grass, herbivores and predators in their movement decisions. As movement affects foraging success and predator-prey encounter rates, it is an important determinant of lifetime reproductive success and, hence, subject to natural selection. Selection pressures are strongly varying in space and time, leading to rapid evolution and rich eco-evolutionary dynamics. Although our model is still very simple, we observe a range of phenomena that do not occur in models with coarser spatial scales and with fewer evolutionary degrees of freedom. The populations in our model do not evolve stable movement patterns but instead exhibit intricate evolutionary dynamics, including regular cycles between movement strategies and the spontaneous advent of novel strategies and counter-strategies. Regularly, stable polymorphisms emerge, where qualitatively different movement strategies coexist for extended periods of time. For example, we observed the evolutionary emergence of sit-and-wait predators, while other predators were chasing their prey. The movement strategies in herbivores and predators determine the spatial pattern of resource depletion and predator-prey encounters, which together induce the emergence of intriguing ecological patterns. Over the generations, these patterns are fluent, as they change with the evolution of the underlying movement strategies. Rapid transitions between patterns (e.g. between spiral waves and static spots) can occur; these reflect coevolutionary changes rather than changes in external conditions. We will now discuss some of these findings in more detail and in the context of other models of predator-prey coevolution.

Previous work on the coevolution of predator-prey movement has also produced interesting patterns, such as evolutionarily optimal strategies leading to ‘predator-prey chases’ across habitats (Abrams 2007) and the coupling of ecological dynamics across habitats via evolving conditional strategies (Flaxman et al. 2011). However, in these studies ‘movement’ corresponds to a choice between a small number of densely populated habitats. In the habitat patches, the interactions of predators and their prey is not modelled at the individual level but by patch-level dynamic equations (but see Patin et al. 2020 for an exception). In contrast to these habitat choice models, we consider movement in a fine-grained spatial environment. From ecological models, it is known that in such fine-grained environments the interplay of predator and prey movement can induce a diversity of spatial ecological patterns, including rotating spirals and static stripes or spots (Hassell et al. 1991, Comins & Hassell 1996, Alonso et al. 2002, Banerjee 2015). Our model shows that, by adding the evolution of movement strategies to the ecological models, various of these patterns emerge in a single simulation run, and that repeated switches between patterns can occur even though the ecological parameters have not changed..

Behavioural polymorphisms commonly occur in our simulations. They can either be fleeting (Supplementary fig. S1) or persistent (fig. 3); they can be relatively stationary (fig. 3B) or strongly fluctuating (fig. 2C); and they can either involve a small number of discrete behavioural types (fig. 3B) or a continuous spectrum of coexisting strategies (fig. 3C). Polymorphisms are also predicted by analytical models of coevolution (Senthilnathan & Gavrilets 2020), but they occur much more frequently in more mechanistic individual-based models (Botero et al. 2010, Long & Weissing 2020). The behavioural dimorphism emerging in the herbivore population in most of our simulations (figs 3 and 4) can be interpreted as the coexistence of two behavioural types that balance predator avoidance and foraging in alternative ways. Behavioural biologists might classify the two types of individuals as bold and shy personalities. From a single-species perspective, it is well known that resource competition and predator-prey interactions can induce behavioural polymorphisms (Quinn & Cresswell 2005, Bell & Sih 2007, Wolf et al. 2007, 2008, Wolf & Weissing 2010). Our simulations show that systematic behavioural variation can also emerge in two species due to coevolution (Fig 4). The existence of behavioural polymorphism can have important ecological and evolutionary implications (Sih et al. 2012, Wolf & Weissing 2012). In predator-prey systems, intraspecific variation can, for example, be crucial for the sustained persistence of both species (Senthilnathan & Gavrilets 2020). This is illustrated by the simulation in Fig. 4 where the breakdown of behavioural polymorphism led to the extinction of the predator species.

Our model produces evolutionary oscillations that are reminiscent of the trait cycles classically found in models of unidimensional traits. These models either consider Mendelian traits, where trait cycles consist in oscillating allele frequencies (population genetics models, Nuismer et al. 2005, Kopp & Gavrilets 2006, Cortez & Weitz 2014), or quantitative traits, where trait cycles are produced by oscillations in the trait value of a monomorphic population (quantitative genetics, Gavrilets 1997, Mougi & Iwasa 2010, Cortez 2018, adaptive dynamics, Dieckmann et al. 1995, Dieckmann & Law 1996, Marrow et al. 1996). A mixture of both approaches, where predators carry a quantitative trait and prey a Mendelian trait, has been shown to produce complex population dynamics such as anti-phase cycles, in-phase cycles and chaotic dynamics (Yamamichi & Ellner 2016). Both frequency-dependent and mutation-limited oscillations naturally occur in our model. In polymorphic populations with a small number of coexisting behavioural morphs (fig. 3), we observed regular oscillations in the frequencies of these morphs. In the absence of behavioural types, other oscillations occurred, which are based on the repeated substitution of traits. These oscillations are mutation-limited in that they depend on the occurrence of specific mutations that can replace the current trait. Due to the stochasticity of the mutation process, the latter trait cycles are much less regular than the oscillations between discrete behavioural types.

We kept our model as simple as possible, in order to demonstrate that not much structure is required for obtaining the eco-evolutionary patterns described above. It would be interesting to study the implications of features such as sexual reproduction and inheritance, variable population sizes at birth, and a spatially heterogeneous resource distribution, but this is beyond the scope of our study. Here we only discuss the implementation of movement strategies by three weighing factors (*w_g_*, *w_h_*, and *w_p_*) in our model. This way, there are three ‘evolutionary degrees of freedom’ in our model, which is considerably more than in conventional predator-prey coevolution models that typically consider a one-dimensional trait space in both the predator and the prey population. Enhancing the degrees of freedom is important, because it allows for much more rapid and unpredictable evolution due to the emergence of ‘surprising’ (and sometimes counterintuitive) strategies and counter-strategies, the advent of novel forms of behaviour, and by allowing for behaviour that is (at least partly) stochastic and unpredictable. We anticipate that, quite generally, models with more evolutionary degrees have much richer eco-evolutionary dynamics than most conventional models.

Having said this, we are fully aware that our behavioural model is still unrealistically simple and well-behaved. Our model could be extended quite naturally by basing the calculation of environmental suitability on a more complex algorithm, such as an evolving artificial neural network (ANN) (Huse et al. 1999, Enquist & Ghirlanda 2005, Morales et al. 2005). Evolving regulatory networks (including ANNs) have a number of important features (Wagner 1979, Van den Berg & Weissing 2015, Van Gestel & Weissing 2016), such as the emergence of cryptic variation (since the same phenotypic strategy can be encoded by very different networks), which allows for much faster evolvability in the face of environmental change. Perhaps most importantly, network models tend to have an intricate genotype-phenotype map, implying that small mutations can have large and unexpected implications at the phenotypic level. It is therefore not surprising that ANN-based pilot simulations on predator-prey coevolution exhibit even richer eco-evolutionary dynamics than occurring in the present model.

## Acknowledgements

We thank P.R. Gupte, R. Bilderbeek and the MARM group at the University of Groningen for valuable discussion, comments, and suggestions. F.J.W. and C.N. acknowledge funding from the European Research Council (ERC Advanced Grant No. 789240).

## Data, code and materials

All data are available on https://dataverse.nl/privateurl.xhtml?token=4cb06f00-15f5-4949-9de5-e0b87f74d1a5. The source code is available under https://github.com/christophnetz/Cinema_git

## Supplementary Material

In this supplement, we give additional information to figures 2,3 and 4. Simulation videos can be accessed in the digital resources. The source code and an executable of the model can be downloaded from GitHub.

### Contents

Figure S1: Evolution of all three weights of prey and predators, compare figure 2

Figure S2: Evolution of all three weights of prey and predators, compare figure 3

Figure S3: Landscape snapshots, compare figure 3

Figure S4: Information usage, compare figure 3

Figure S5: Weight evolution of all weights of prey and predators, compare figure 4 Digital resources: https://dataverse.nl/privateurl.xhtml?token=4cb06f00-15f5-4949-9de5-e0b87f74d1a5

Simulation videos 1-3: Landscape dynamics in replicate simulations of figure 2. Predator-prey ratio 1:1; offspring dispersal radius 1.

Simulation video 4: Landscape dynamics in replicate simulations of figure 3. Predator-prey ratio 1:10; offspring dispersal radius 10.

Source Code: https://github.com/christophnetz/Cinema_git

### Supplementary information to figure 2

This simulation was conducted with a predator-prey ratio of 1:1 under local offspring dispersal with a radius of 1 (for other default parameters see model description). Replicate simulation movies can be viewed [here]. The landscape dynamics reflect the ecological interactions which in turn are shaped by the evolution of movement decision mechanisms, encoded by the weights w_g_, w_h_ and w_p_ (see equation 1 in the main text). The evolution of all six weights, three per herbivore and predator population each, is depicted in Supplementary figure S1. Notice that there are distinct periods with similar weight configurations, between which rapid shifts occur. For example, from generation 10,000 to 18,000, the herbivore population contains a dimorphism with regard to the predator density weight w_p_, similar to the more stable polymorphism we observe in figure 3. The occurrence and vanishing of the positive-valued morph is associated with a distinct phase of performance and ecological pattern shown in fig. 2B.2. From generation 6,000 to 10,000, the predator population even contains three morphs with respect to the weights for grass and herbivore density, w_g_ and w_h_. One of these morphs is positive for both weights, the second is effectively neutral, whilst the third and least-frequent morph is negative for both. Again this population composition is tightly associated with a performance phase and the ecological pattern in fig. 2B.1. As the third morph goes extinct around generation 10,000, performance and ecological pattern changes. Towards the end of the simulation, herbivores evolve a rather constant strategy consisting in a strongly positive grass density weight w_g_, a strongly negative herbivore density weight w_h_, and a strongly positive predator density weight w_p_. However, this state, corresponding to the ecological pattern shown in fig. 2B.5, is punctuated by the repeated occurrence of herbivores with a negative predator-density weight that escape the surrounding predators and spread across the landscape (fig.2B.6). Whenever this morph arises, we observe a peak in herbivore performance. This morph cannot establish itself permanently, though: Its agglomerations collapse under the pressure of advancing predators as individuals move away from predators. The morph with a positive herbivore density weight on the other hand does not move away from predators and can therefore form stable agglomerations. It is worth noting that these results do not necessarily transfer to empirical systems, as individuals in reality of course rarely move towards predators. Yet, this result demonstrates the creativity, with which antagonists exploit each other’s strategic weaknesses created by phenotype and external constraints (in this case given by our model, otherwise given by the laws of nature), and the far-reaching ecological consequences this can have.

### Supplementary information to figure 3

We next considered a predator-prey ratio of 1:10 and local offspring dispersal with a radius of 10. Under these conditions the landscape is much less structured than in the previous simulation (see Supplementary fig. S3, movie) and the predator-prey ratio is closer to general empirical expectations. The herbivore population evolves a dimorphism for the predator density weight w_p_ (see figure 3). While one morph exhibits a negative-valued weight w_p_ and therefore avoids predators, the other is weakly positive-valued and predator-prone. This counter-intuitive strategy can be understood when looking at the evolution of all six weights that is depicted in Supplementary figure S2: The predator evolved a negative predator-density weight, and a (slightly) positive predator-density weight therefore makes the movements of that herbivore morph less predictable to the predator. The other two weights of the predator show a positive preference for grass and herbivore density. Thus the predators chase prey, while at the same time expecting to find herbivores more often in areas of high grass. The other two weights of the herbivore population show a preference for grass, but an avoidance of herbivore densities. This could be an adaptation to avoid intraspecific competition for resources as well as heightened predation risk in the presence of conspecifics.

The weight values together form the movement strategy of an individual, but it is still difficult to assess the overall importance of the three environmental cues: grass density, herbivore density and predator density follow different distributions, and therefore influence the movement decisions to different extends even if all three weights were the same. We could derive the distributions from the overall landscape, but this would ignore the heterogeneous distribution of individuals and the bias introduced by selective movement. Thus to correct for this, we produced summary statistics of the cues experienced by all individuals of a population during a single generation. We can then multiply weight values (here we can segregate by morph) with the mean absolute deviation of grass density, herbivore density and predator density cues, giving us the information usage during each generation (Supplementary fig. 4). In the beginning of the simulation, predators as well as w_p_-positive and w_p_-negative herbivore morphs move primarily according to grass density. The w_p_-positive herbivore morph considers herbivore densities second, while considering predator densities least, and maintains this profile of information usage throughout the simulated time. The w_p_-negative herbivore morph on the other hand undergoes a gradual shift between generations 3,000 and 5,000 towards considering predator densities more strongly. After generation 5,000, this morph moves primarily according to predator densities, with grass density and herbivore density ranking second and third. The predator population follows this development in a rapid shift around generation 6,000, such that it then matches the information usage profile of the w_p_-negative herbivore morph. These results demonstrate how difficult it can be to determine the general importance of predator densities, herbivore densities and resource densities. Movement strategies can be heterogeneous within populations and change dynamically through time.

### Supplementary information to figure 4

Figure 4 shows a special case in which predators were only allowed to sense their own presence, with a predator-prey ratio of 1:10 and local offspring dispersal with a radius of 10. We chose to show this special case, because it illustrates two types of trait cycles and evolutionary-induced extinction. For completeness, figure S5 shows the all 4 weights. One can see that the grass and herbivore density weights of the herbivore populations hitchhike on the oscillations induced by the interaction between the predator density weights in predators and herbivores. However, the initial genetic variation is depleted during the transitions indicated by t1, t2, t3 and t4.

### Simulation Code and Availability

Our model program implements a visual representation of the simulated landscape, in which users can visually inspect pattern formation and even the movements and interactions of single individuals. We encourage the reader to watch the video clips of our simulations that we uploaded here [https://dataverse.nl/privateurl.xhtml?token=4cb06f00-15f5-4949-9de5-e0b87f74d1a5], as they confer a much more detailed impression of the eco-evolutionary dynamics than the landscape snapshots (fig. 2, Supplementary fig. S) Everyone who would like to experiment themselves with our model finds our model code on GitHub, [https://github.com/christophnetz/Cinema_git]. The program produces a graphical output of the landscape each time step (as shown in videos 1-4), a heatmap of the weight distributions as well as a performance plot. Users can also pause the simulation and zoom in on single individuals. Please refer to the readme file for more detailed instructions, but note that the program depends on C++ libraries that are only available for the Windows operating system.

---

**Figure S1:**
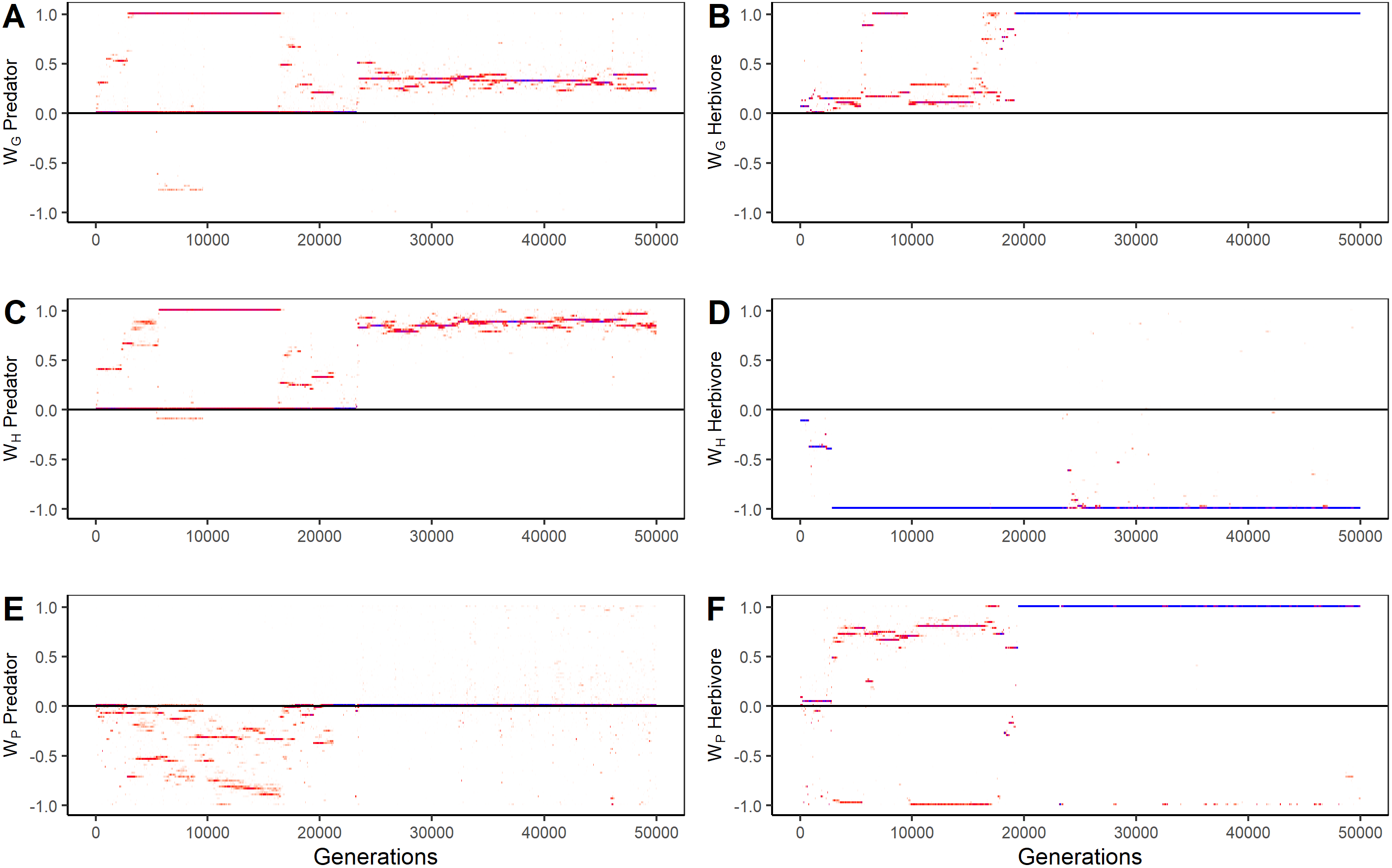
Weight evolution of herbivores and predators, compare figure 2. Frequencies go from 0.0 (=white) to 0.3 (=red) and 1 (=blue). Weight values were multiplied with 100 and transformed by a hyperbolic tangent. Predator-prey ratio 1:1; offspring dispersal radius 1.

**Figure S2:**
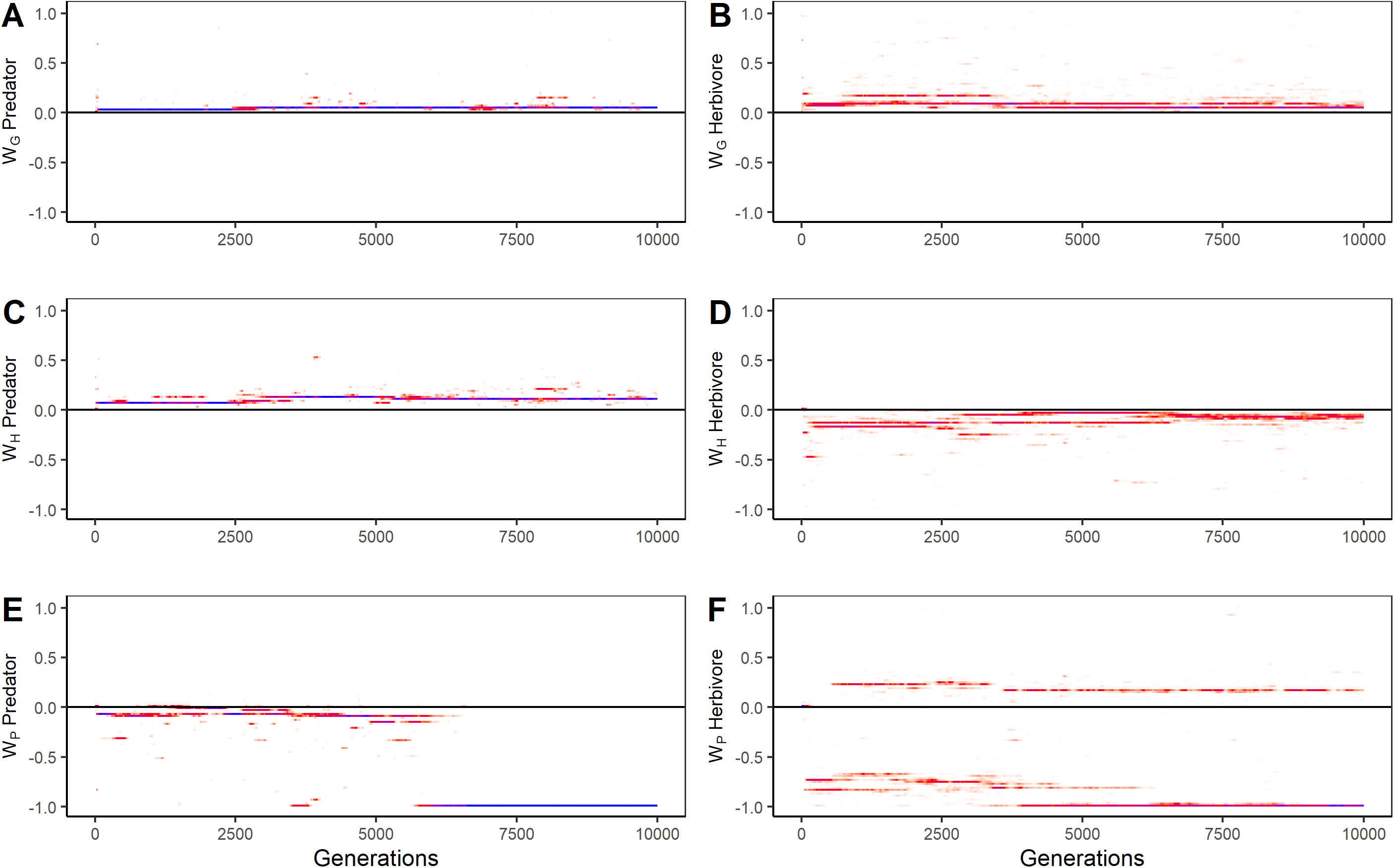
Weight evolution of herbivores and predators, compare figure 3. Frequencies go from 0.0 (=white) to 0.3 (=red) and 1 (=blue). Weight values were multiplied with 100 and transformed by a hyperbolic tangent. Predator-prey ratio 1:10; offspring dispersal radius 10.

**Figure S3:**
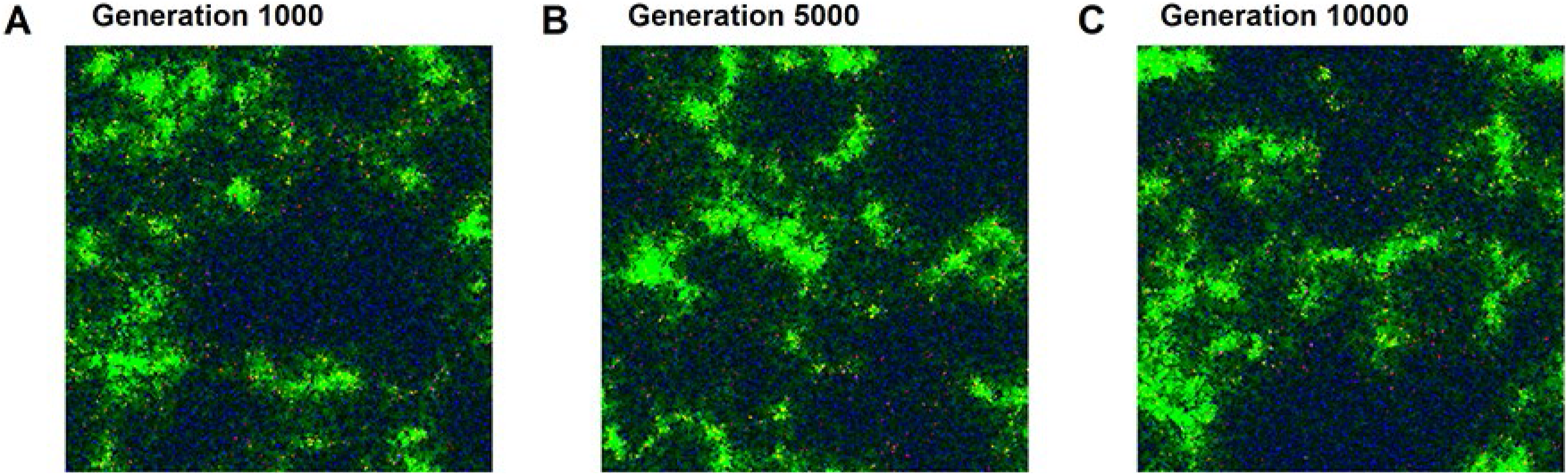
Landscape snapshots of a replicate simulation of figure 3 at generations 1000, 5000 and 10000. Grass density is shown in green, herbivore density in blue and predator density in red. Other colors emerge from additive color mixing, yellow for example signifies areas of high grass and predator density, purple that herbivores and predators occupy the same area. Predator-prey ratio 1:10; offspring dispersal radius 10.

**Figure S4:**
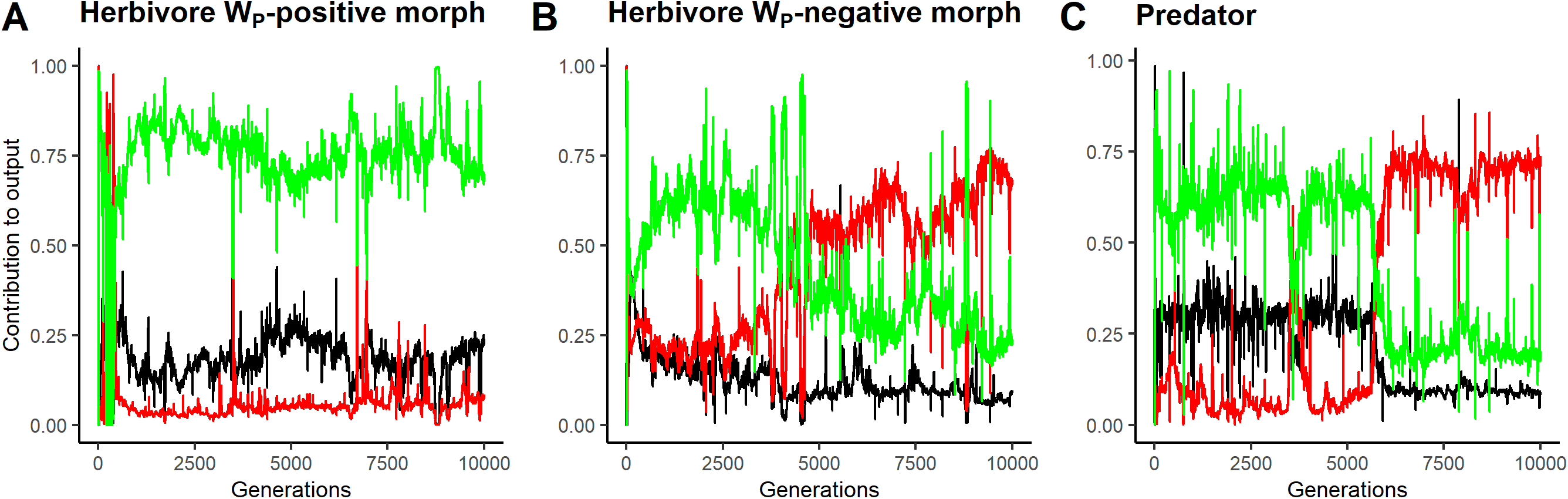
Average information usage of the predator-prone herbivore morph (w_p_-positive morph), the predator-avoiding herbivore morph (w_p_-negative morph), and the predator population, compare figure 3. Predator-prey ratio 1:10; offspring dispersal radius 10.

**Figure S5:**
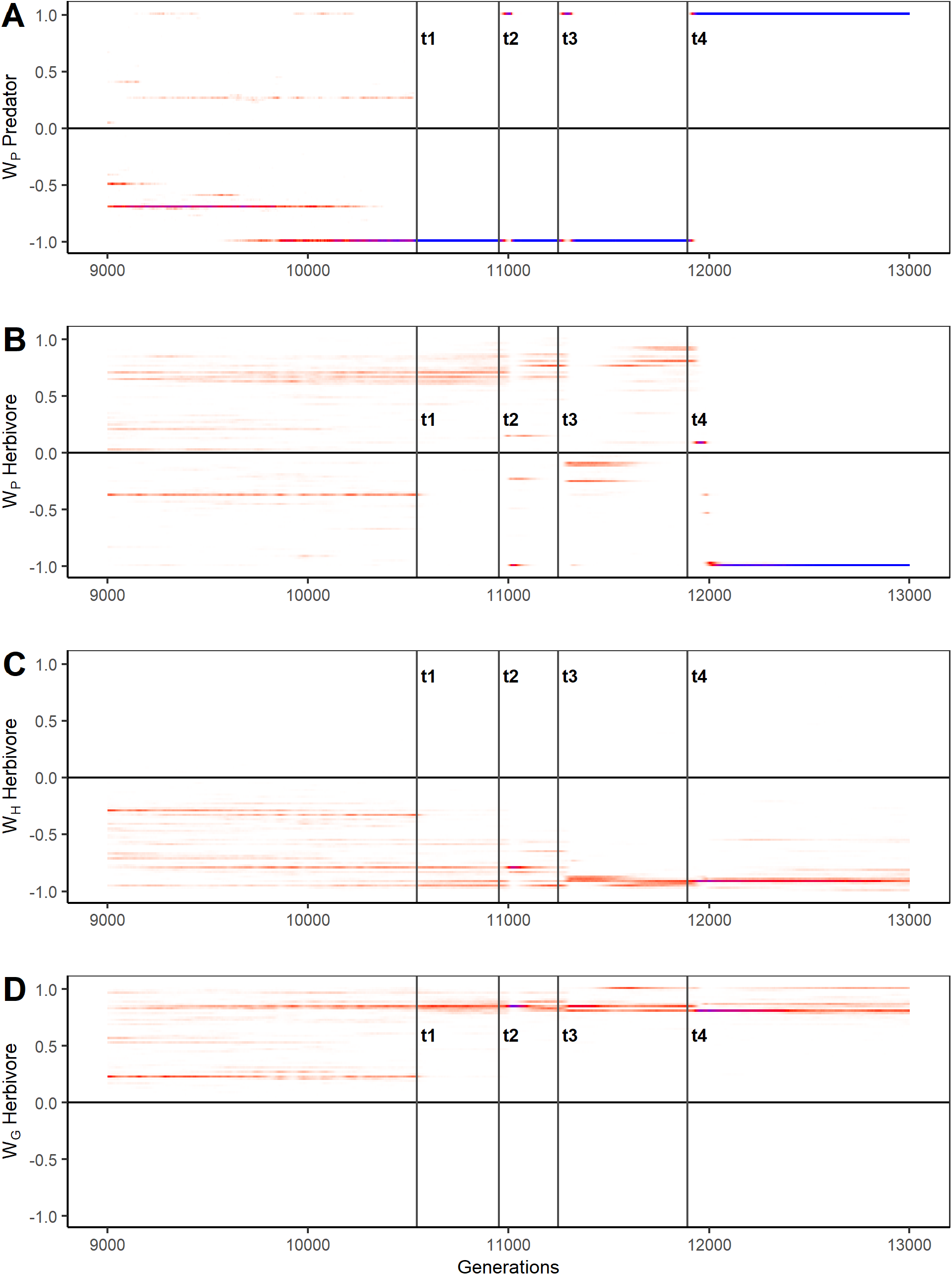
Cut-out from a simulation in which predators could only sense other predators. Evolution of all three weights of herbivores and the single predator weight, compare figure 4. Frequencies go from 0.0 (=white) to 0.3 (=red) and 1 (=blue). Weight values were multiplied with 100 and transformed by a hyperbolic tangent. Predator-prey ratio 1:10; offspring dispersal radius 10. t1 marks the extinction of the predator-prone predator morph at generation 10544, t2, t3 and t4 the reintroduction of this morph via mutation at generations 10953, 11249 and 11891.

